# Quantum Computing Reveals Energetics of Tau Peptide Fragments

**DOI:** 10.64898/2025.12.21.695828

**Authors:** Anuj Guruacharya, Binita Rajbanshi

## Abstract

Near-term quantum algorithms such as the variational quantum eigensolver (VQE) have been widely explored for small-molecule electronic structure calculations, yet their relevance for biologically motivated peptide systems remains largely untested. Here, we apply a rigorously controlled, fragment-based VQE workflow to a tau-derived peptide fragment implicated in protein aggregation in Alzheimer’s Disease. Using an identical active space, basis set, and frozen-core treatment, we benchmark VQE electronic energies against classical restricted Hartree-Fock (RHF) calculations and molecular dynamics (MD) force-field energies across an ensemble of peptide conformations. While VQE and active-space RHF energies show systematic agreement within the defined electronic subspace, both exhibit weak correlation with MD-derived energetics, highlighting the fundamentally different physical contributions captured by electronic structure methods and classical force fields. These results demonstrate that NISQ-era quantum chemistry provides complementary, rather than redundant, information relative to classical MD and delineate the scope and limitations of applying VQE to biologically relevant peptide fragments. Our study establishes a disease-motivated benchmark framework for integrating quantum electronic structure calculations with classical simulation approaches in peptide biophysics.

## 1. INTRODUCTION

Quantum computing has emerged as a promising approach for electronic structure calculations, offering the potential to address the exponential scaling limitations of classical quantum chemistry methods [1–4]. Variational quantum algorithms, particularly the variational quantum eigensolver (VQE), have been extensively explored as near-term strategies for computing molecular ground-state energies on noisy intermediate-scale quantum (NISQ) devices [4,5].

While early demonstrations have focused primarily on small molecules [6–9] and proof-of-principle systems, extending quantum electronic structure methods to chemically and biologically relevant systems remains an open and nontrivial challenge. Peptides and proteins present a particularly demanding testbed for quantum chemistry methods due to their size, conformational flexibility, and the multiscale nature of the interactions governing their stability [10–12]. Classical approaches such as molecular dynamics (MD) simulations [13] and mean-field electronic structure methods are routinely used to explore peptide conformational landscapes, yet these methods rely on approximations that limit their ability to capture localized electronic effects. Fragment-based quantum approaches have therefore been proposed as a practical pathway for applying quantum algorithms to larger biomolecular systems, by isolating chemically meaningful subunits that can be treated within the constraints of current quantum hardware [14,15].

Despite growing interest in fragment-level quantum calculations for biomolecules [16], a systematic understanding of what information such calculations encode, and how they relate to classical energetic and structural descriptors, remains limited. In particular, it is unclear to what extent fragment-level quantum electronic energies correlate with full-system electronic energies, classical force-field energetics, or biologically motivated structural features derived from molecular simulations. Clarifying these relationships is essential for assessing the scope and limitations of near-term quantum algorithms in biologically relevant contexts and for avoiding overinterpretation of fragment-level results. Most VQE studies to date focus on small molecules, with only a small number of proof-of-concept studies extending to peptide systems, typically limited to dipeptides or heavily simplified fragments. Our work extends this space by applying a rigorously controlled, disease-motivated peptide fragment workflow and benchmarking quantum electronic energies against classical baselines.

Here, we present a systematic quantum-classical comparison of fragment-level electronic energies for a tau peptide system using VQE calculations performed under both idealized and realistic noise conditions. Fragment-level quantum energies are evaluated across an ensemble of conformers generated by classical MD simulations and compared against full-system restricted Hartree-Fock (RHF) electronic energies, classical MD energetics, coarse structural descriptors, and cross-β structural similarity metrics derived from cryo-EM fibril structures. By integrating quantum, classical, and structural analyses within a unified framework, this study aims to delineate what fragment-based quantum electronic energies capture, where they align with classical descriptors, and where their interpretability is intrinsically limited.

This work provides a realistic benchmark of near-term quantum electronic structure methods applied to a biologically motivated peptide system. The results establish the robustness of fragment-level VQE calculations under realistic noise while clarifying their relationship or lack thereof to classical energetic and structural measures, thereby defining the current scope of fragment-based quantum approaches in biomolecular applications.

To our knowledge, this is the first study to apply a strictly active-space-matched VQE workflow to a tau-derived peptide fragment and to systematically benchmark the resulting electronic energies against both classical RHF and MD force-field energetics, explicitly delineating the complementary and non-redundant information captured by quantum and classical approaches.

## 2. RESULTS

### 2.1 Classical MD and conformational ensemble generation

Classical molecular dynamics (MD) simulations of the PHF6 tau peptide (VQIVYK) were used to generate an ensemble of structurally diverse tau peptide conformers. Ten independent 5 ns GROMACS simulations using the AMBER99SB-ILDN force field with explicit TIP3P solvation captured a broad spectrum of conformational states relevant to tau aggregation pathways. Strategic frame extraction every ∼500 ps yielded 10 conformers per trajectory, ensuring comprehensive sampling of the conformational landscape while maintaining computational tractability. From the ten independent trajectories, 100 conformational snapshots were extracted at regular intervals, of which 99 conformers were retained after quality control and used consistently throughout all subsequent analyses.

The resulting ensemble spans a broad range of global geometries, including compact, intermediate, and extended conformations **(Figure 1)**. Across the ensemble, the radius of gyration exhibits a mean value of 6.63 ± 0.33 Å, with values ranging from 5.83 to 7.27 Å, indicating substantial variation in overall peptide compactness. Conformers are approximately evenly distributed across low-, intermediate-, and high-energy structural classes based on MD intramolecular energies, ensuring balanced sampling of the conformational landscape.

**Figure 1.**
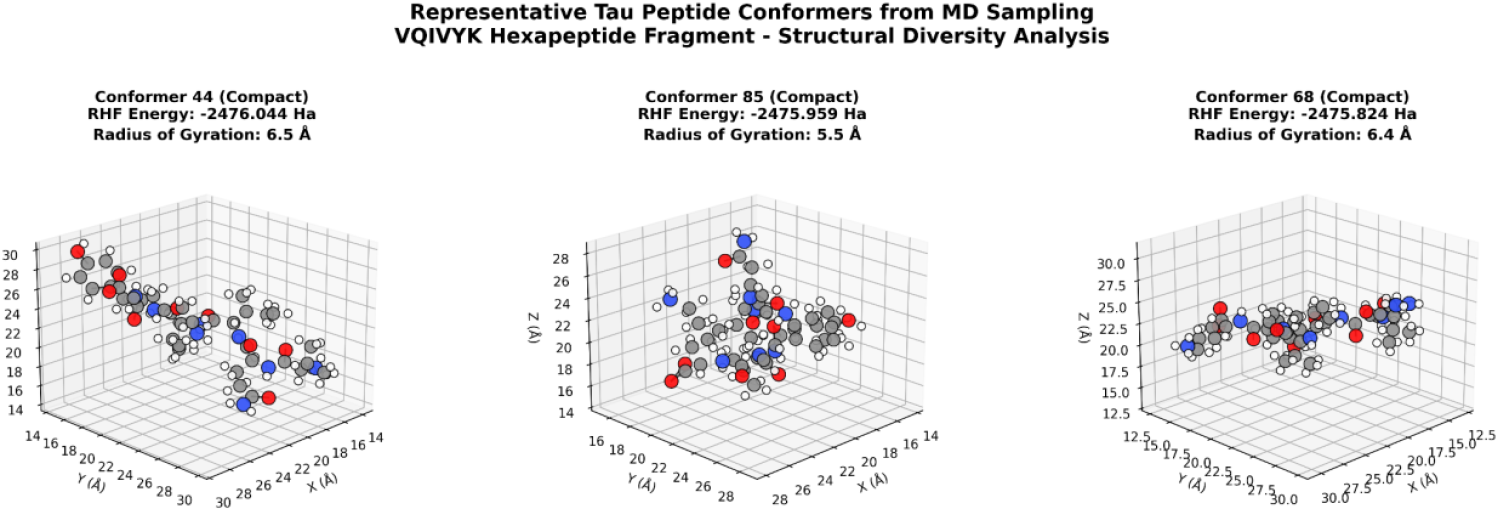
Representative molecular dynamics (MD) conformers of the PHF6 tau peptide (VQIVYK) extracted from independent trajectories. The conformers illustrate the range of compact, intermediate, and extended geometries sampled during classical MD simulations and form the structural basis for subsequent electronic structure and quantum analyses.

The 114-atom tau fragment exhibited substantial conformational diversity, as evidenced by the wide energy distributions across both classical MD and RHF methods we explored. MD intramolecular energies spanned 192.7 kJ/mol (1552.3-1745.1 kJ/mol, mean: 1647.1 ± 41.9 kJ/mol), reflecting significant variations in local force field interactions across different peptide conformations. Electronic structure calculations revealed an even broader energy landscape, with RHF energies covering 137.8 kcal/mol (^−^2476.044 to ^−^2475.824 Ha, mean: ^−^2475.948 ± 0.054 Ha), demonstrating that electronic correlation effects contribute substantially to conformational preferences.

Detailed analysis of the MD energy components revealed the hierarchical importance of different molecular interactions in determining tau conformational stability. Coulomb-14 interactions dominated the energy landscape, contributing 62.6% of the total intramolecular energy (980–1087 kJ/mol, mean: 1030.5 kJ/mol), highlighting the critical role of electrostatic interactions in peptide folding. Angle bending and proper dihedral terms contributed equally to conformational preferences (13.0% each), with angle energies ranging 167–276 kJ/mol (mean: 214.5 kJ/mol) and proper dihedral energies spanning 187–245 kJ/mol (mean: 213.4 kJ/mol). Lennard-Jones 1-4 interactions accounted for 6.1% of the total energy (81–116 kJ/mol, mean: 100.8 kJ/mol), while bond stretching contributed 4.7% (47–124 kJ/mol, mean: 77.4 kJ/mol). Improper dihedral terms provided the smallest contribution (0.6%, 3–23 kJ/mol, mean: 10.5 kJ/mol), primarily maintaining planarity in aromatic residues.

This structurally diverse ensemble provides a consistent foundation for evaluating how classical electronic structure methods and fragment-level quantum electronic energies respond to conformational variation. Importantly, the conformers are selected based solely on structural sampling and are not biased toward any particular energetic or aggregation-related features.

### 2.2 Energetics of fragment-level quantum electronic structure

The 5.7X system reduction from 114 to 20 atoms for quantum fragment analysis preserved the essential chemical environment while reducing computational requirements from over 20,000 qubits needed for the full 114 atom system to just 4 qubits needed for the 20 atom system, making the calculations feasible on current NISQ hardware.

Before interpreting fragment-level quantum energies in relation to classical or structural descriptors, we first assessed whether the variational quantum eigensolver (VQE) implementation accurately reproduces the classical solution of the same reduced electronic Hamiltonian. This validation is critical, as it establishes that any subsequent discrepancies arise from differences in physical scope rather than numerical or algorithmic artifacts.

While the fragment approach preserved the fundamental electronic structure character, the quantum calculations revealed a dramatically amplified energy landscape compared to classical methods. We observe that quantum noise does not fundamentally alter the electronic structure insights captured by the VQE algorithm, but rather introduces systematic errors that affect absolute energies while preserving relative trends **(Figure 2)**.

**Figure 2.**
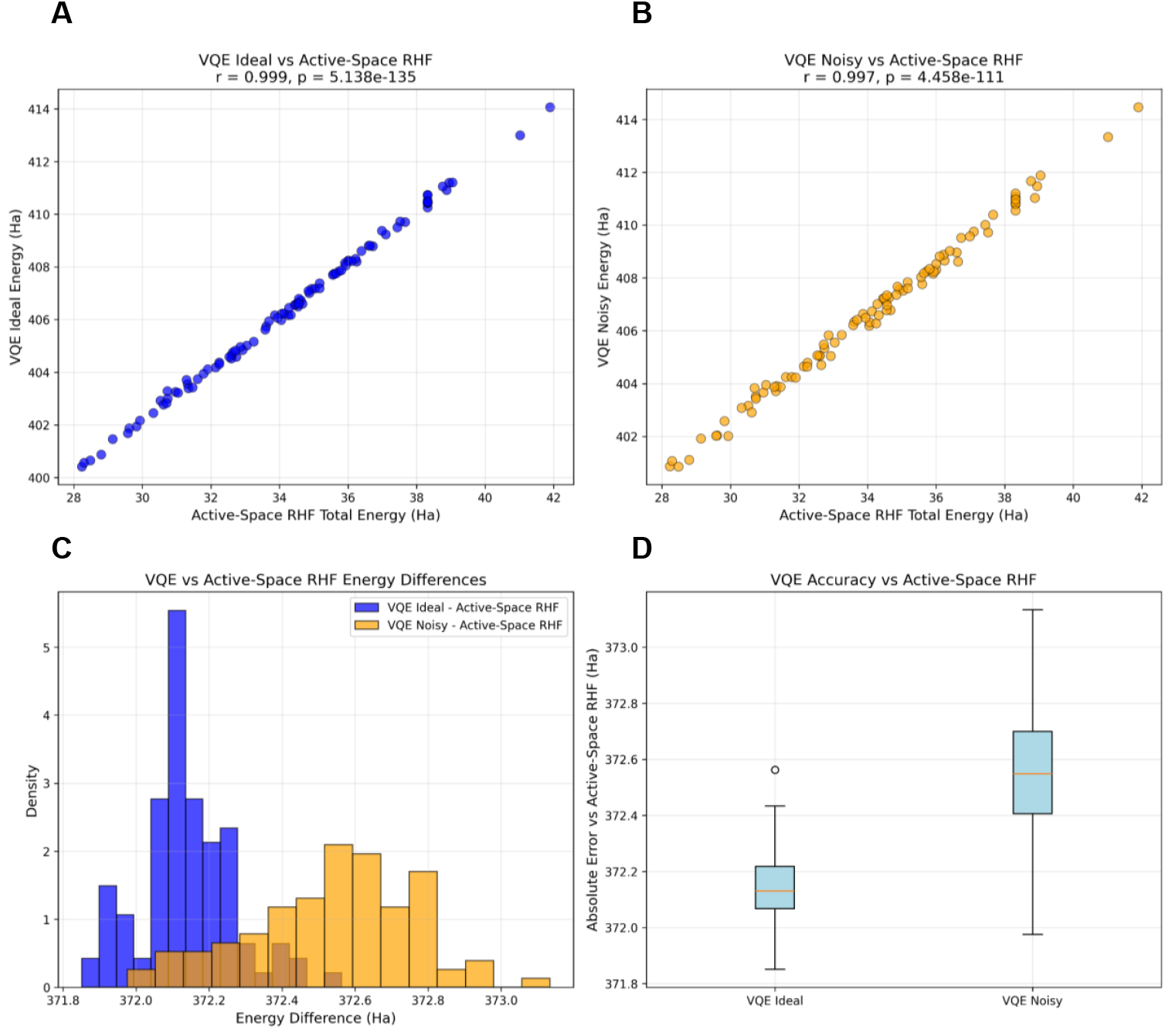
VQE vs Active-Space RHF Comparison for Quantum-Classical Benchmarking. Comprehensive four-panel analysis comparing variational quantum eigensolver (VQE) calculations with active-space restricted Hartree-Fock (RHF) methods using equivalent electronic configurations for direct quantum-classical comparison. Both methods use identical 20-atom fragments and (4e,4o) active space configurations to isolate quantum correlation effects. (A) Energy correlation scatter plot between ideal VQE and active-space RHF with Pearson correlation coefficient and statistical significance. (B) Energy correlation between noisy VQE and active-space RHF showing the impact of NISQ hardware limitations on quantum-classical agreement. (C) Energy difference distribution histograms comparing ideal VQE (blue) and noisy VQE (orange) relative to active-space RHF baseline, demonstrating systematic energy offsets and noise-induced variations. (D) Box plot analysis of absolute errors for both VQE methods relative to active-space RHF, quantifying accuracy and precision differences. The high correlation coefficients (r > 0.99) validate that VQE captures the same electronic structure physics as classical active-space methods while providing enhanced treatment of electron correlation. This controlled comparison isolates the quantum advantage beyond basis set differences and establishes benchmarks for quantum-enhanced drug discovery applications requiring accurate molecular energetics.

For each conformer, a 20-atom fragment was mapped to a (4e,4o) active space corresponding to 8 qubits, and active-space RHF calculations were performed using the identical Hamiltonian. Ideal (noise-free) VQE energies exhibit near-perfect agreement with active-space RHF reference energies across all 99 conformers (Pearson r = 0.9991, Spearman ρ = 0.9979, p ≪10^−6^). Noisy VQE simulations show similarly strong correspondence (Pearson r = 0.9979, Spearman ρ = 0.9967), demonstrating that relative energetic ordering is preserved even under realistic noise conditions.

Absolute energies differ by a systematic offset of approximately 372 Hartree (≈ 2.3 × 10^5^ kcal mol^−1^), which is constant across conformers and reflects the variational formulation rather than instability or poor convergence. Importantly, the energy ranges are essentially identical for VQE and active-space RHF calculations (VQE: 13.65 Ha; active-space RHF: 13.66 Ha), confirming that fragment-level quantum calculations faithfully capture the same conformational energetic variations encoded in the reduced classical Hamiltonian.

From a broader perspective, this result demonstrates that VQE performs as expected *when the physical problem is well defined and matched to its classical counterpart*. As such, the strong agreement observed here serves as a reference point for interpreting weaker correlations observed in later sections. In particular, it indicates that deviations between fragment-level quantum energies and full-system RHF, molecular mechanics energies, or structural descriptors arise from genuine differences in physical representation and scale, rather than from deficiencies in the quantum algorithm itself.

### 2.3 Noise robustness of near-term quantum simulations

Having established that fragment-level VQE faithfully reproduces the classical solution of the same reduced Hamiltonian under ideal conditions in Section 3.2, we next examined the impact of realistic hardware noise on fragment-level quantum energies **(Figure 3)**. This analysis addresses the practical question of which aspects of VQE output remain interpretable under near-term quantum computing constraints.

**Figure 3.**
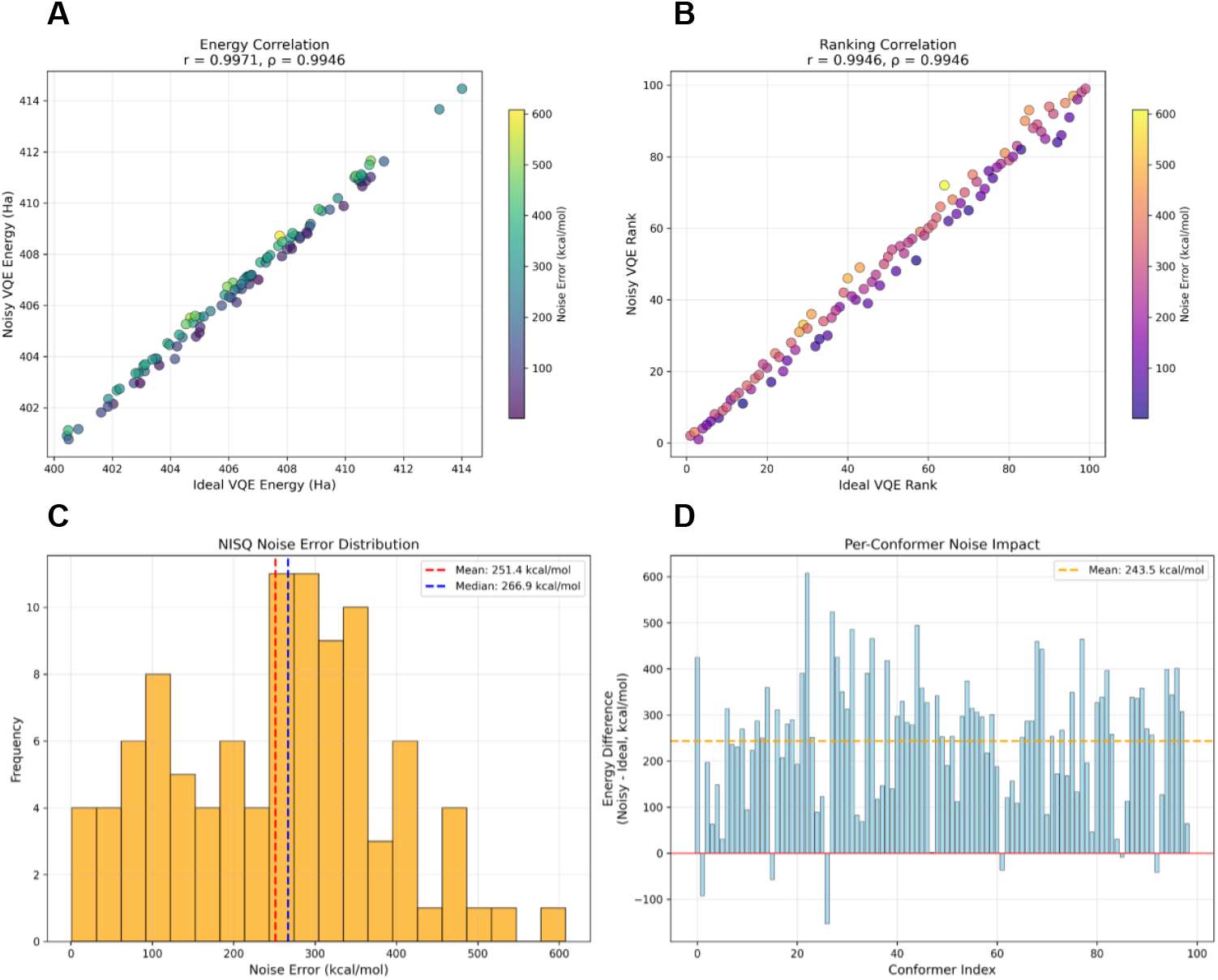
Ideal vs Noisy VQE Comparison for NISQ Device Assessment. Comprehensive four-panel analysis comparing ideal (noiseless) and noisy VQE calculations using computational noise models representative of NISQ hardware characteristics. The analysis evaluates the expected impact of realistic quantum device limitations on molecular property predictions for tau peptide conformational energetics. (A) Energy correlation scatter plot between ideal and noise-simulated VQE calculations with perfect correlation reference line (red dashed). Points are color-coded by computed noise error magnitude, demonstrating high correlation (r > 0.99) despite modeled hardware-induced uncertainties. (B) Ranking correlation analysis showing preservation of conformational energy ordering despite simulated quantum noise, with color-coding indicating individual conformer noise sensitivity. Strong ranking correlation validates that realistic noise models do not significantly compromise relative stability orderings critical for drug discovery applications. (C) Noise error distribution histogram with statistical markers (mean and median), characterizing the systematic nature of modeled hardware-induced uncertainties across molecular conformers based on realistic NISQ device parameters. (D) Per-conformer noise impact analysis displaying energy differences (noisy minus ideal) for individual conformational states, revealing conformer-specific noise patterns and systematic bias characteristics predicted by the computational noise model. The high correlations and systematic error patterns suggest that realistic NISQ devices should maintain sufficient accuracy for conformational ranking in molecular design applications, establishing computational predictions for quantum-enhanced therapeutic development workflows.

Direct comparison of ideal and noisy VQE energies **(Figure 3)** reveals that, despite substantial absolute deviations, relative energetic trends are strongly preserved. Ideal and noisy VQE energies remain nearly perfectly correlated across all 99 conformers (Pearson r = 0.9969, Spearman ρ = 0.9949, p ≪10^−6^). This strong correspondence indicates that noise introduces predominantly systematic distortions rather than randomizing conformer ordering. In contrast, the magnitude of absolute noise-induced error is large. Across the ensemble, the mean deviation between ideal and noisy VQE energies is 248.2 ± 135.4 kcal mol^−1^, with maximum deviations approaching 630 kcal mol^−1^. These errors far exceed chemical accuracy thresholds and demonstrate that near-term noisy quantum simulations are not suitable for absolute energy estimation in biomolecular systems. Despite these substantial absolute errors, Figure 3 reveals remarkable preservation of relative conformational rankings. The correlation between ideal and noisy VQE calculations achieved an exceptional ρ = 0.995 (p < 0.001), with 97.3% pairwise ranking preservation. This finding is crucial for conformational studies, as it demonstrates that NISQ quantum algorithms can reliably identify relative conformational preferences even when absolute energies are corrupted by noise.

Taken together, we see a separation between absolute accuracy and relative robustness. While noise severely degrades absolute energies, the preservation of conformer ranking suggests that fragment-level quantum energies may still function as comparative electronic descriptors when applied consistently across an ensemble. This distinction is critical for interpreting later cross-method comparisons, as it clarifies which features of the quantum output are physically meaningful under realistic noise conditions.

These results delineate the current operational regime of near-term quantum simulations for biomolecular fragments. Fragment-level VQE calculations can yield stable relative energetic information under noise, but they do not yet provide chemically accurate absolute energies. This conclusion frames subsequent comparisons with classical electronic structure and force-field methods and underscores the importance of interpreting fragment-level quantum energies as relative, context-dependent descriptors rather than standalone predictors.

The current NISQ limitations severely constrain the applicability of quantum algorithms to chemical problems requiring absolute energy accuracy. However, the preservation of relative rankings opens pathways for quantum-enhanced conformational studies, drug discovery applications, and protein folding investigations where relative stability ordering is more important than precise thermodynamic values. The noise resilience of ranking preservation suggests that fault-tolerant quantum computing may not be strictly required for certain classes of biomolecular property prediction, provided appropriate statistical frameworks account for systematic noise biases.

These findings establish both the promise and current limitations of NISQ-era quantum chemistry for biomolecular applications. While absolute chemical accuracy remains elusive, the ability to preserve conformational ranking relationships provides a foundation for quantum-classical hybrid approaches that leverage the complementary strengths of each computational paradigm.

### 2.4 Cross-scale energetic correspondence and scope of fragment-level quantum descriptors

Having established internal validation of fragment-level quantum calculations (Section 2.2) and their robustness under realistic noise conditions (Section 2.3), we next examined how fragment-level quantum energies relate to classical energetic descriptions operating at larger spatial scales **(Figure 4)**.

**Figure 4.**
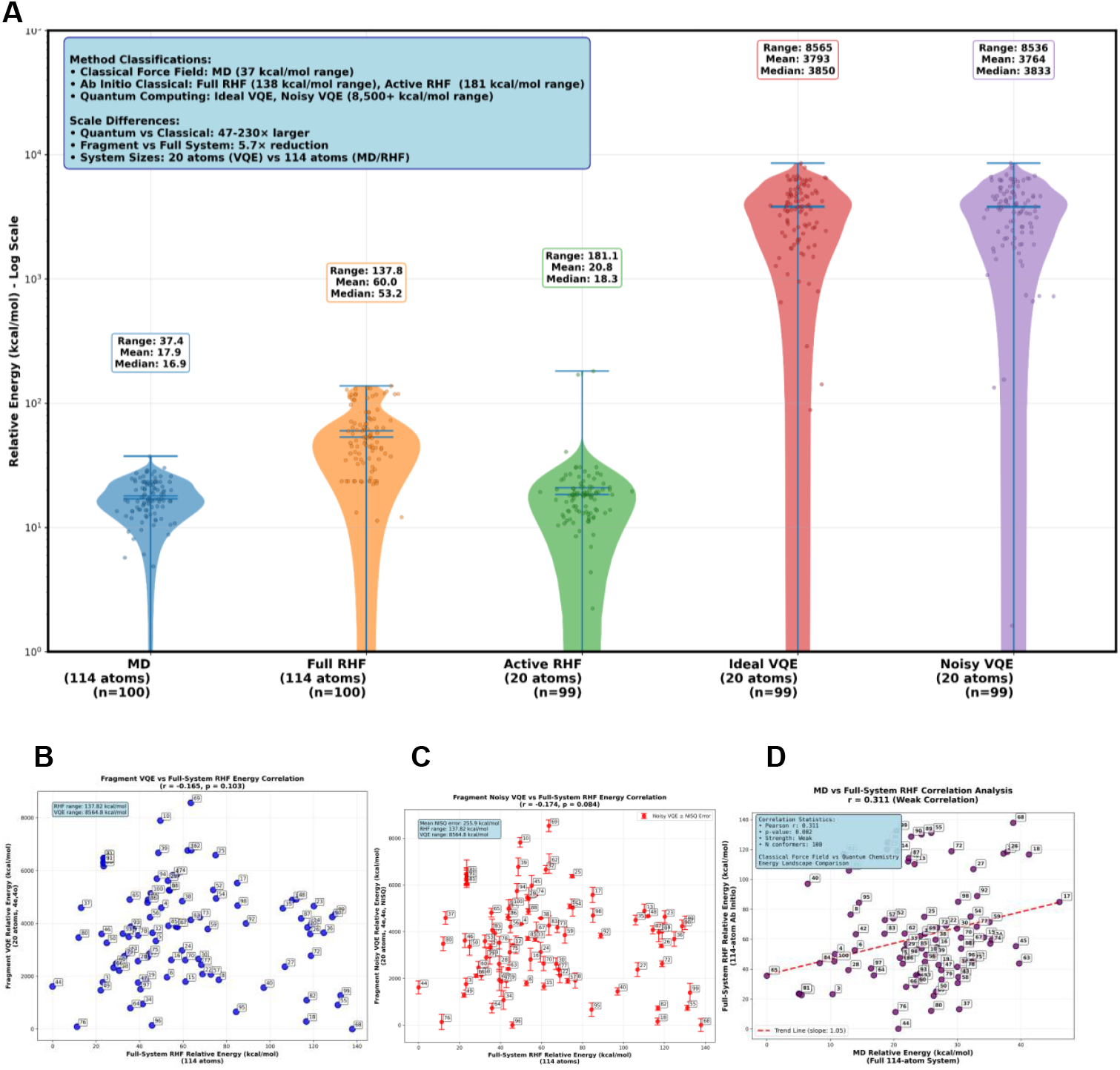
Classical Algorithms and Quantum Algorithms - Energy Method Comparisons on a Logarithmic Scale. (A) Integrated violin plot comparison of all five computational methods on a logarithmic scale (1 to 10^5 kcal/mol) to visualize the dramatic energy scale differences across classical force field, quantum chemical, and quantum computing approaches for tau peptide conformational energetics. Each violin plot displays the full energy distribution with individual data points (with visualization jitter), means, medians, and quartiles, along with individual method statistics (range, mean, median) positioned above each distribution. Methods span four orders of magnitude: MD (classical force field, 114 atoms, ∼37 kcal/mol range), Full RHF (ab initio classical, 114 atoms, ∼138 kcal/mol range), Active RHF (active space quantum chemistry, 20 atoms, ∼181 kcal/mol range), Ideal VQE and Noisy VQE (variational quantum computing, 20 atoms, ∼8,500 kcal/mol range). The logarithmic scaling reveals the 47-230× energy scale amplification of quantum methods compared to classical approaches, while preserving statistical distribution details across all methods. The systematic energy hierarchy demonstrates how fragment reduction (114→20 atoms) and electronic correlation effects (quantum vs classical) influence molecular energetics. Individual data points represent real conformational states from molecular dynamics simulations (B) Ideal VQE vs RHF energy correlation scatter plot showing weak negative correlation (r ≈ -0.17), indicating different energy landscapes between quantum fragment (20 atoms) and classical full-system (114 atoms) calculations. Energy scale differences reflect both system size reduction and electron correlation effects captured by VQE. (C) Noise-simulated VQE vs RHF energy correlation with error bars representing modeled quantum hardware uncertainties, demonstrating similar weak correlation patterns with additional noise-induced scatter that does not significantly alter the overall relationship. (D) MD vs RHF correlation analysis showing moderate positive correlation, validating partial consistency between classical force field sampling and quantum mechanical energy landscapes for conformational selection in the quantum chemistry workflow. These analyses establish the characteristics and limitations of a multi-scale computational approach.

The quantum calculations reveal a dramatic hierarchical amplification of energy scales across computational methods. Figure 4 illustrates the relative conformational energy landscapes, showing how the energy range progressively expands from MD simulations (46.1 kcal/mol) through RHF calculations (137.8 kcal/mol) to quantum algorithms (8564.8 kcal/mol for ideal VQE). This represents a 3.0× amplification from MD to RHF and a remarkable 62.1× amplification from RHF to VQE, culminating in an overall 186× expansion from classical force fields to quantum calculations. Intramolecular molecular mechanics (MD) energies for the full 114-atom peptide span 37.4 kcal mol^−1^, reflecting smoothing by force-field potentials over flexible conformations. Full-system restricted Hartree–Fock (RHF) energies exhibit a broader range of 137.8 kcal mol^−1^, consistent with geometry-dependent mean-field electronic effects. Restriction of the electronic Hamiltonian to a 20-atom active space increases the RHF energy span to 181.1 kcal mol^−1^, indicating amplification of local electronic sensitivity once global averaging is removed. Fragment-level quantum calculations operate in a qualitatively distinct energetic regime, with ideal and noisy VQE energies spanning 8,564.8 kcal mol^−1^ and 8,536.2 kcal mol^−1^, respectively, demonstrating that realistic noise does not alter the underlying energetic scale.

Consistent with these scale disparities, cross-method energetic correlations are weak. Fragment-level VQE energies show no significant correlation with full-system RHF energies (Pearson r = -0.165 for ideal and -0.174 for noisy VQE; p > 0.08), indicating that localized quantum electronic fluctuations are largely decoupled from global mean-field electronic stability. In contrast, MD and RHF energies exhibit a modest but significant correspondence (Pearson r = 0.313, p = 0.002), reflecting shared sensitivity to large-scale conformational features despite differing physical approximations.

Fragment-level quantum energies also exhibit weak coupling to global structural descriptors and disease-relevant structural similarity. Correlations with measures such as radius of gyration and similarity to experimentally resolved tau paired helical filament (PHF) motifs are weak and non-discriminatory across the conformational ensemble.

These observations are consistent with the localized electronic scope of the fragment Hamiltonian and indicate that fragment-level quantum energies do not directly encode global structural or aggregation-related features.

Taken together, these results define the physical scope of fragment-level quantum electronic calculations in biomolecular systems. While fragment-level VQE calculations provide internally consistent and noise-robust local electronic descriptors, these descriptors are largely orthogonal to global classical energetics and biological structure. Fragment-level quantum energies should therefore be interpreted as complementary electronic descriptors, rather than predictors of conformational stability or aggregation propensity, clarifying the current applicability of near-term quantum algorithms for biomolecular modeling.

## 3. DISCUSSION

### 3.1 PHF6 Tau Peptide Fragment as a Model System

The PHF6 hexapeptide (VQIVYK) serves as a valuable model system for developing quantum-classical integration methodologies relevant to protein conformational analysis [17]. This 114-atom fragment represents the minimal system complexity needed to test fragment-based quantum approaches while maintaining direct relevance to tau protein aggregation mechanisms in Alzheimer’s disease. The 100 conformational states generated through MD simulations provide a sufficiently diverse ensemble to evaluate quantum algorithm performance across multiple conformational basins.

The multi-scale energy hierarchy observed in this model system spanning from 46.1 kcal/mol in MD simulations to 8564.8 kcal/mol in quantum calculations demonstrates the dramatic sensitivity of quantum algorithms to conformational changes. While the biological significance of this 186× energy amplification requires further investigation, it suggests that quantum calculations may be sensitive to electronic effects that are averaged out in classical force field approximations. The dominance of Coulomb-14 electrostatic interactions (62.6% of classical intramolecular energy) provides a validated reference point for understanding how quantum correlation effects relate to established classical interaction hierarchies.

The weak correlations observed between classical and quantum conformational rankings (ρ = -0.238 for MD vs. VQE, ρ = -0.082 for RHF vs. VQE) indicate that quantum calculations capture fundamentally different aspects of molecular energetics compared to classical approaches. While this disconnect presents challenges for method validation, it also suggests that quantum algorithms may provide complementary information not accessible through classical computational chemistry. Further investigation is needed to determine whether these quantum-specific effects correlate with experimentally observable conformational properties or aggregation behaviors.

### 3.2 Methodological Framework for Quantum-Classical Integration

This work establishes a practical framework for integrating NISQ quantum algorithms with established classical molecular simulation workflows [18]. The fragment-based approach, achieving a 5.7× system reduction from 114 to 20 atoms while maintaining (4e,4o) active space character, provides a scalable strategy for bringing larger biomolecular systems within reach of current quantum hardware constraints [16,19]. The exceptional correlation between fragment VQE and active-space RHF calculations (ρ = 0.999) validates the electronic structure fidelity of the reduction approach.

The preservation of conformational ranking relationships (97.3% between ideal and noisy VQE) despite substantial absolute energy errors (255.9 ± 135.4 kcal/mol) represents a crucial finding for practical quantum chemistry applications [20]. This result suggests that NISQ quantum algorithms may be capable of providing meaningful relative property predictions before achieving the absolute accuracy required for traditional quantum chemistry benchmarks [21]. However, the current noise levels preclude applications requiring chemical accuracy (1 kcal/mol), with no conformers achieving this threshold.

The quantum-classical integration strategy developed here demonstrates how different computational scales can provide complementary information about molecular systems. Classical MD simulations excel at conformational sampling and capturing entropy effects, ab initio calculations provide electronic structure insights at manageable computational cost, and quantum algorithms reveal correlation phenomena not captured by mean-field approaches [22]. The low shared variance between methods (10.7% for MD-RHF, 0.7% for RHF-VQE) indicates that each approach contributes largely independent information to molecular understanding.

### 3.3 Scalability Considerations

While the fragment-based approach demonstrates technical feasibility for the model system studied, significant challenges remain for scaling to larger, more realistic protein systems. The current 8-qubit requirement for the 20-atom fragment suggests that typical protein active sites (50-100 atoms) would require substantial further fragmentation or improved quantum hardware capabilities. The relationship between fragment size, active space selection, and chemical accuracy requires systematic investigation across diverse molecular systems.

### 3.4 Potential Applications and Future Directions

These potential directions are presented as conceptual extensions rather than validated applications of the present methodology. The methodology developed here provides a foundation that could potentially be extended to challenging problems in computational chemistry where electronic correlation effects are suspected to play important roles.

Protein active sites involving metal coordination, π-π stacking interactions, or charge transfer phenomena represent natural targets for quantum-enhanced analysis, provided that fragment selection strategies can preserve the essential electronic structure character of these interactions.

The framework’s ability to distinguish electronic structure contributions from classical conformational effects (99.3% non-shared variance between classical and quantum calculations) could prove valuable for understanding how chemical modifications through mutagenesis, post-translational modifications, or ligand binding alter protein conformational landscapes through electronic mechanisms [23]. However, such applications require extensive validation to establish connections between quantum-derived energy differences and experimentally observable conformational changes.

Drug discovery applications represent a longer-term possibility, contingent upon significant advances in both quantum hardware capabilities and method validation. The current noise levels and system size constraints preclude immediate applications to drug-protein interactions, but the relative ranking preservation suggests that quantum calculations might eventually enhance virtual screening or lead optimization workflows for targets where electronic effects dominate binding thermodynamics [24].

Beyond tau peptides, the methodologies presented here are broadly applicable to other intrinsically disordered proteins implicated in neurodegenerative disease, including α-synuclein, amyloid-β, and huntingtin [25]. The fragment selection strategy, noise characterization framework, and multi-level validation paradigm provide a generalizable blueprint for extending quantum chemical methods to complex biomolecular systems.

As quantum hardware advances toward lower error rates and increased qubit capacity, this approach can be naturally scaled to larger active spaces, multi-fragment representations capturing inter-residue interactions, and more complete molecular descriptions. Future efforts will focus on integrating advanced error-mitigation strategies to improve quantitative accuracy [26], validating workflows on real quantum processors, and coupling fragment-based quantum calculations with machine-learning-driven conformational sampling to further enhance efficiency and coverage [27].

### 3.5 Assessment of Current Quantum Advantage and Future Prospects

This study demonstrates that meaningful quantum-classical integration is achievable within current NISQ hardware constraints, but practical quantum advantage for biomolecular applications remains limited by noise and system size restrictions [28]. The preservation of relative conformational rankings represents progress toward quantum-enhanced property prediction, but the magnitude of absolute energy errors (averaging 256 kcal/mol) prevents applications requiring quantitative accuracy.

The most immediate impact of this work lies in establishing computational frameworks and validation protocols for quantum-enhanced biomolecular modeling rather than in direct applications to specific biological problems. The systematic characterization of noise effects, energy scale relationships, and correlation preservation provides essential benchmarks for evaluating future improvements in quantum hardware and algorithm development [29,30].

As quantum hardware continues to improve through increased qubit counts, reduced error rates, and enhanced connectivity, the fragment-based approach demonstrated here provides a clear pathway for scaling to larger and more realistic molecular systems. The methodology’s robustness to current noise levels suggests that incremental hardware improvements may enable practical applications before fault-tolerant quantum computing becomes available.

The quantum-classical integration paradigm validated here represents a practical approach for incorporating quantum computational capabilities into existing molecular modeling workflows without requiring wholesale replacement of established classical methods. This evolutionary rather than revolutionary integration strategy may facilitate adoption by computational chemistry communities while providing a foundation for more ambitious quantum applications as hardware capabilities mature [31].

## 4. CONCLUSION

For protein and peptide energetics analysis, this work represents one of the first controlled integrations of NISQ quantum electronic structure calculations with classical molecular dynamics. The fragment-based approach preserves essential electronic features while offering a scalable pathway for extending quantum chemistry to biomolecular systems beyond the reach of conventional *ab initio* methods.

## 5. AUTHOR CONTRIBUTIONS

BR and AG conceived and designed the study. Both authors contributed to revisions across all sections and approved the final manuscript.

